# Harnessing Transformers to Generate Protein Sequences Prone to Liquid Liquid Phase Separation

**DOI:** 10.1101/2024.03.02.583105

**Authors:** Abdul Wasim, Ushasi Pramanik, Anirban Das, Pikaso Latua, Jai S. Rudra, Jagannath Mondal

## Abstract

Understanding the molecular grammar that governs protein phase separation is essential for advancements in bioinformatics and protein engineering. This study leverages Generative Pre-trained Transformer (GPT)-based Protein Language Models (PLMs) to decode the complex grammar of proteins prone to liquid-liquid phase separation (LLPS). We trained three distinct GPT models on datasets comprising amino acid sequences with varying LLPS propensities: highly predisposed (LLPS+ GPT), moderate (LLPS-GPT), and resistant (PDB* GPT). As training progressed, the LLPS-prone model began to learn embeddings that were distinct from those in LLPS-resistant sequences. These models generated 18,000 protein sequences ranging from 20 to 200 amino acids, which exhibited low similarity to known sequences in the SwissProt database. Statistical analysis revealed subtle but significant differences in amino acid occurrence probabilities between sequences from LLPS-prone and LLPS-resistant models, suggesting distinct molecular grammar underlying their phase separation abilities. Notably, sequences from LLPS+ GPT showed fewer aromatic residues and a higher fraction of charge decoration. Short peptides (20-25 amino acids) generated from LLPS+ GPT underwent computational and wet-lab validation, demonstrating their ability to form phase-separated states in vitro. The generated sequences enriched the existing database and enabled the development of a robust classifier that accurately distinguishes LLPS-prone from non-LLPS sequences. This research marks a significant advancement in using computational models to explore and engineer the vast protein sequence space associated with LLPS-prone proteins.

## I. INTRODUCTION

Biomolecular condensates, formed through liquid–liquid phase separation (LLPS), play crucial roles in various cellular functions within the cytoplasm and nucleus. Understanding the contributions of the molecular interactions that drive LLPS is key to grasping the fundamentals of these assemblies and exploring their potential applications[1–4].

Emerging investigations over last decade on drivers of LLPS has particularly noted the predisposition of proteins which are intrinsically disordered. Intrinsically disordered proteins (IDPs) represent a large subset of proteome[5] lacking a well-defined secondary structure. They possess the ability to transition between various folded conformations based on the specific function they are required to perform. The currenly accepted hypothesis posits that IDPs, prior to aggregation, might experience spontaneous liquid-liquid phase separation (LLPS). Experimental observations indicate that IDPs can undergo LLPS in vitro when present in a solution above specific concentrations and under particular environmental conditions. Consequently, the LLPS of various proteins in diverse environments—whether physiological or otherwise—is currently under investigation through a combination of experimental[6–11], computational[12–20], and analytical[21–26] approaches.

To unravel the distinctive features of IDPs stemming from their amino acid sequences, extensive analysis of the primary sequences of numerous IDPs has been conducted[27, 28]. Furthermore, studies have employed various metrics derived solely from the primary structures of different protein classes to accurately quantify and characterize the LLPS-forming properties of IDPs[28, 29]. Collectively, these investigations suggest the existence of different set of rules governing the amino acid arrangement of IDPs and folded proteins. Consequently, these rules, often referred to as the “molecular grammar” of these proteins, contribute to shaping their unique phenotypes. The development of models, such as the “sticker and spacer” model[30], has been instrumental in comprehending the LLPS-forming behavior of IDPs. Such rules have introduced empirical views such as presence of low complexity regions (LCRs), that are enriched in polar, aromatic and proline and glycine residues, as potential drivers of LLPS. However, recent investigations[31–33] have also showed that numerous exception exists where deviation from these traits can readily lead to LLPS and the fact that microenvironments potentially would play a very important role.

Together these investigations enrich but complicate the so called “molecular grammar” of proteins that are essential for LLPS. The smaller size of existing LLPS-prone protein-sequence datasets also does not help the cause. In this study, we set out to understand if we can make use of generative artificial intelligence (AI) based approaches that can objectively learn the traits present in pre-curated protein sequences which are distinct in their LLPS propensity and harness the learning for generation of new sequences. For this purpose, here we employ AI based Protein Language Models (PLMs). PLMs use blocks of transformer neural networks to understand the context in a given protein sequence and hence can not only be used to understand the inherent set of rules or “molecular grammar” present in the sequence but also can be used to generate new sequences by traversing the protein sequence space.

The advent of Transformers[34] and the availability of robust computing hardware have led to the developement of powerful large language models (LLMs). One notable example is the renowned ChatGPT[35], which has captivated both experts and non-experts with its capabilities. The remarkable accuracy achieved by current LLMs can be attributed to the utilization of transformer neural networks. Transformers excel in comprehending sequential data by leveraging token and positional embeddings to discern the “context” within a sequence. With an ample amount of data and models of sufficient scale (comprising billions of parameters), transformers exhibit a proficiency in understanding language in a probabilistic manner. This proficiency grants them an implicit understanding of the inherent set of rules or grammar governing the language. Since the amino acid sequences of proteins adhere to a set of rules very specific to proteins and entirely different to natural languages, language models developed for protein sequences have been coined as Protein Language Models (PLMs) as they understand the grammar of the amino acid sequences rather than a language.

In this investigation, we employ decoder-only transformer-based neural networks to train Generative Pre-trained Transformer (GPT) models on protein sequences, aiming to grasp the intrinsic molecular grammar of these sequences. Utilizing pre-curated protein datasets—LLPS+, LLPS- and PDB*[27] we train individual GPT models on each dataset. Subsequently, we generate a substantial number of amino acid sequences with varying lengths using these models. Our initial analysis involves examining and comparing specific features of these generated sequences with corresponding features extracted from the original datasets. Furthermore, we analyse an amino acid occurrence probability matrix to gain insights into the molecular grammar of proteins which are or are not LLPS prone. We find that the difference in the occurrence probabilities of amino-acids in sequences obtained from LLPS-prone (LLPS+, LLPS-) and LLPS-resistant (PDB*) GPT models is subtle but sufficient for resulting in differences in phase separation behaviour. To validate our findings, we derive a subset of short (20-25 amino acid long) peptides from LLPS+ GPT for computational and in vitro validation. For instance, via molecular dynamics simulation we predict that a short protein derived from the LLPS+ GPT model can undergo spontaneous liquid-liquid phase separation (LLPS) displaying a finite upper critical solution temperature (UCST). We then follow it up in wet lab where these peptide sequences are found to undergo LLPS in ambient condition. Finally we use the GPT models to build a balanced dataset for the purpose of training a tree-based ML classifier for distinction between LLPS prone sequences and sequences that do not undergo LLPS..

## II. RESULTS

### A. Training GPT Models on Amino Acid Sequences

In this study we use a special type of Machine Learning architecture called “Transformers”[34] to generate three distinct Protein Language Models (PLMs). Transformers have been used in the context of Natural Language Models, such as ChatGPT[35] or LLAMA[36] with great success. This led to the exploration of Transformers for understanding the molecular grammar of proteins. Notable works are ProstT5[37] and ProtBERT[38]. However, most of the PLMs developed till now are “encoder” based PLMs and can be used to encode amino acid sequences into vectors, with the information of the order of the amino acids intact. In this study we trained “decoder” based PLMs, characterized Masked attention instead of simple attention. We used the datasets reported by Saar *et al*.[27]: LLPS+, LLPS-, and PDB* for training three PLMs, each corresponding to one dataset.

The LLPS+ dataset comprises 137 unique protein sequences, each capable of undergoing LLPS *in-vitro* under ambient conditions and having critical concentrations to induce LLPS at most 100 *µ*M. The PLM trained on LLPS+ has been named LLPS+ GPT.

The LLPS-dataset has only 82 entries, each capable of undergoing LLPS but under more drastic conditions. They either require higher protein concentrations (*>*100 *µ*M) or environments that favor LLPS induction. This PLM has been named LLPS-GPT.

Finally, we also trained a PLM on the PDB* dataset, which has 1562 unique protein sequences, each of which are folded proteins and will not undergo LLPS unless extremely drastic conditions are applied. The corresponding PLM has been named PDB* GPT.

Figure 1 shows an overview of PLM architecture along with the information flow. We shall discuss the information flow and model training later in this section.

**FIG. 1.**
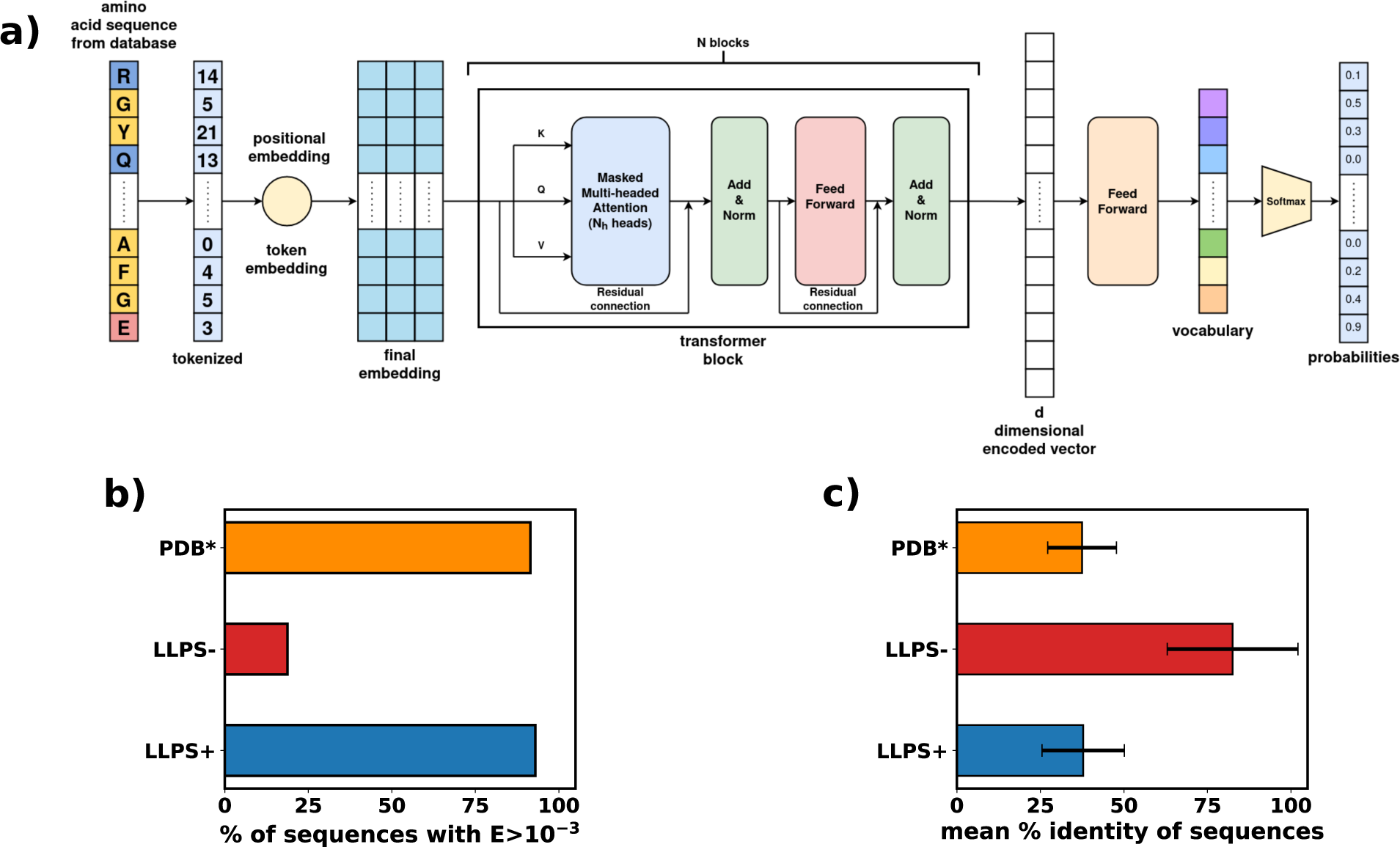
**a)** The architecture of the Transformer based Protein Language Models (PLMs) used and schematic of their training procedure. **b)** Percentage of GPT-generated sequences having E-value greater than 0.001 when compared against the SwissProt database. **c)** Percentage identity of the GPT-generated sequences with sequences present in the SwissProt database

To train each PLM, we first built our vocabulary from the respective dataset. Protein sequences were denoted using the single letter codes for amino acids (see details in*Methods* section). The vocabulary helps us convert the amino acid sequences, which are text, into sequences of integers, which are numbers that can be fed to the neural network. This process is called tokenization (Figure 1). Each token, which is an integer, is then converted into a vector of dimension *d_embed_*. We have used *d_embed_* = 256. This vector is the “token embedding”. Coupled with “positional embedding”, which preserves the information regarding the order of the tokens (see *Methods*), we obtain the final embedding (Figure 1).

Traditionally, token and positional embedding were generated via human-made rules [34]. However, a more generic way is to let the models figure out their own token and positional embedding, which we have implemented here via the “Embedding” module of PyTorch. It should be noted that since the positional embedding depends on the sequence length, the maximum length (*l_max_*) that can be passed to the model is also fixed during model building. In this study, we have used *l_max_* = 128. Hence, a sequence that is being passed into our models, in a single shot, cannot have more than 128 amino acids in it. However, one can generate very long sequences using the models and we shall discuss this later in this section. The final embedding thus will have the dimensions (*l, d_embed_*), where *l* (*≤ l_max_*) is the sequence length fed into the network. *d_embed_* is a model hyperparameter and is fixed during model building.

The final embedding is then passed into *N* consecutive blocks of transformers, with each transformer block processing the data using *N_h_* heads (Figure 1). All our models have been made using *N* = 8 and *N_h_* = 8. Each head looks at the data independently. At the end of the transformer block, a mean of the outputs from each of the heads is passed onto the next block. The output from each transformer block has the shape (*l, d*) where *d* is the model hidden dimension (Figure 1). We have used *d* = 256 for all PLMs we have built. This is then projected onto a vector that has the size of the vocabulary via a feed-forward neural network. A “sigmoid” activation function over the final vector provides the probability of each token/vocabulary to appear next in the sequence (Figure 1). The most probable token is selected as the next amino acid.

For training, sequences of length *l_max_* + 1 are selected. The first *l_max_* amino acids are tokenized and passed into the network. The network tries to predict the last amino acid at the *l_max_* + 1 position. Using a cross-entropy loss and an AdamW optimizer [39], which are routinely used for training such networks, the model is trained iteratively to minimize the error of the loss function (Figure S1). Thus, with each round of training, the model gets better at predicting the next amino acid by learning the rules that govern the order of the amino acids. In other words, it learns the molecular grammar inherent in the set of amino acid sequences that have been used during training. However, the grammar that it learns is inscrutable to us since it is hidden inside very large dimensional weights and biases matrices in each head inside each transformer block.

For the prediction of new sequences, one can start with a single, randomly chosen amino acid. The model will output another amino acid, which can then be appended to the input. This updated input is fed back into the model to predict the next amino acid. By repeating this process in an auto-regressive manner, it is possible to generate long sequences of amino acids (for details see *Methods*). For easy usage of the trained PLMs for new sequence generation, we have created a Google Colab notebook which can be accessed from https://bit.ly/4clObMP.

### B. Distinctiveness of Generated Sequences Compared to Known Proteins

We employed the three trained models (LLPS+ GPT, LLPS-GPT and PDB* GPT) to produce 6000 amino-acid sequences from each model, with lengths ranging from 20 to 200 amino acids (refer to *Methods*). Supporting data files S1-S3 contain the sequences generated using LLPS+ GPT, LLPS-GPT and PDB* GPT respectively in FASTA format. To validate the distinctiveness of the generated sequences from those utilized during training, we initially conducted a multiple sequence alignment using BLAST against the datasets employed for model training. For each sequence we focus on two metrics provided by BLAST that denotes how similar a given sequence is to other sequences present in the a database of choice: E and percentage identity. E determines the statistical significance of a match between two sequences. Higher E represents a random match, i.e. the match is not statistically significant and vice versa. Percentage identity quantifies the residue-level similarity between two sequences. A higher similarity means a more perfect match and vice-versa. Our analysis revealed a no alignment between the generated sequences and those existing in the training datasets. This substantiates that the sequences generated by the model differ significantly from those encountered during training.

Subsequently, we conducted multiple sequence alignments against the SwissProt database for all 6000 sequences. We observed that 99%, 20% and 98% of the sequences generated from LLPS+, LLPS- and PDB* GPT models, respectively, have E-values greater than 0.001 (Figure 1b). This suggests that for LLPS+ and PDB*, almost all of the sequences do not align well with of the already known proteins curated in the SwissProt database. We think that for LLPS-, the dataset coverage of the protein space might have been too small for the LLPS-GPT to be able to generate novel sequences and hence the lower dissimilarity percentage. A similar result is also being shown by the statistics for the percentage identity of the alignments (Figure 1c).

Analyzing the multiple sequence alignment results against SwissProt, a notable observation emerges. The majority of protein sequences generated from the LLPS+ and PDB* models exhibit a noteworthy degree of uniqueness, evident in their E-value and percentage identity statistics. Conversely, sequences generated from the LLPS-model display a higher alignment with proteins already documented in the SwissProt database. It is crucial to emphasize that, during the training phase, our models were never exposed to the SwissProt database. This underscores the remarkable capability of the Protein Language Models (PLMs) trained on protein sequences. Not only do these models adeptly predict entirely novel and unseen proteins, but they also navigate the intricate protein sequence space with remarkable efficiency. This affirms the models’ capacity to generate diverse and distinctive protein sequences, showcasing their potential to contribute meaningfully to the exploration of uncharted areas within the realm of protein sequences.

### C. Learning Embeddings for Different Protein Types During Training

As discussed earlier, embeddings are an integral part of a PLM. In the models that we have developed, the embeddings are learned during training. This implores us to ask the following questions: 1) Are the embeddings for all the PLMs similar when they have not yet been trained? 2) During training, do the embeddings also pick up the differences in the sequences used for their training? 3) Do the PLMs finally learn very different embeddings based on the dataset used to train them?

To answer these questions, we must consider that embeddings are high-dimensional data. To make sense of this data, we project the embeddings onto a two-dimensional “abstract” coordinate space. This coordinate space is a non-linear combination of the original embedding dimensions achieved via training an AutoEncoder (Figure 2a and *SI Methods*).

**FIG. 2.**
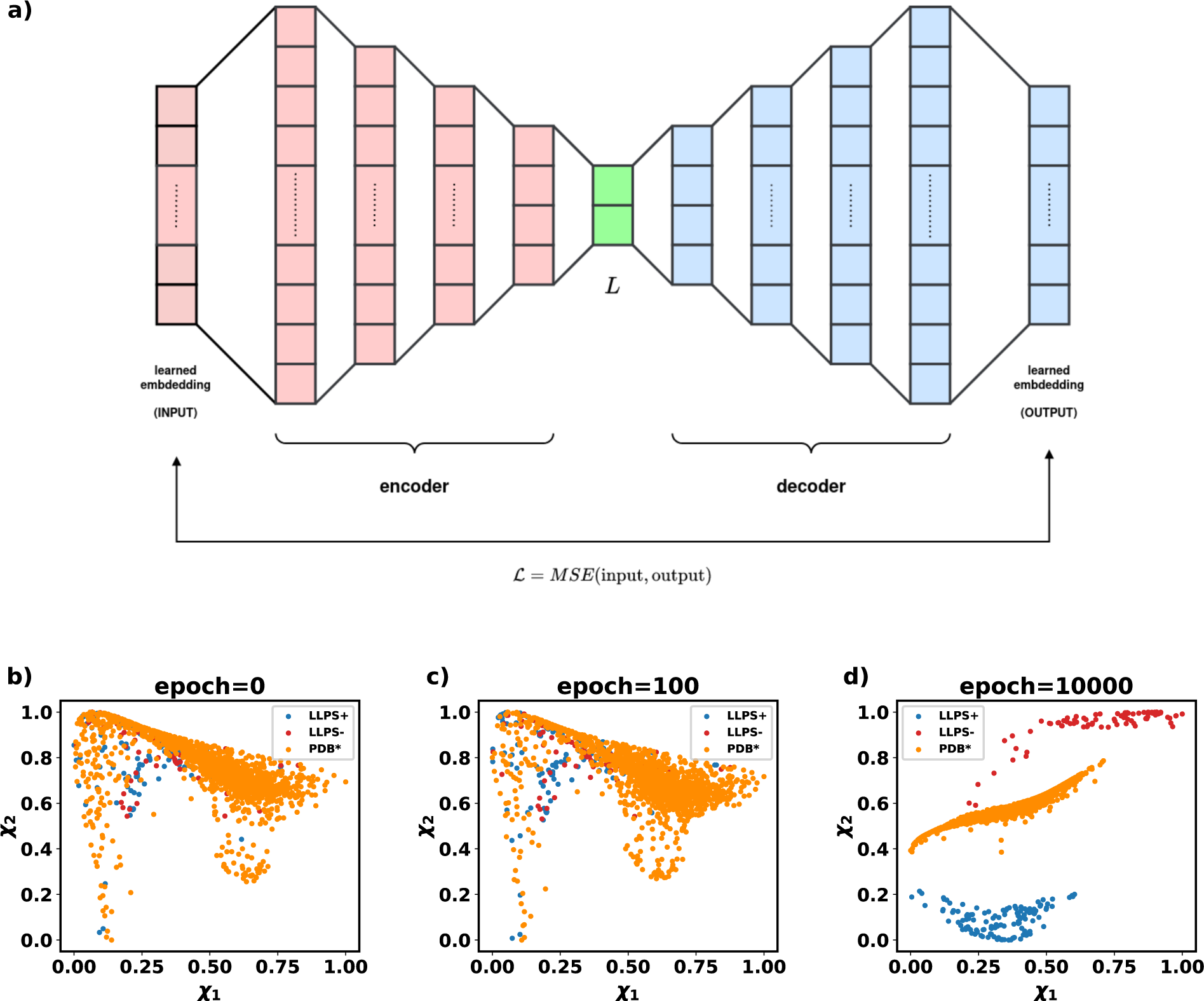
**a)** The architecture of an AutoEncoder. **b)** LLPS+, LLPS- and PDB* embeddings projected onto a latent space at 0th epoch. **c)** LLPS+, LLPS- and PDB* embeddings projected onto a latent space at 100th epoch. **d)** LLPS+, LLPS- and PDB* embeddings projected onto a latent space at 10000th epoch.

An AutoEncoder is a self-supervised algorithm that compresses input data into latent dimensions via the encoder and subsequently decodes the latent dimensions back to the original data using the decoder. The training process aims to minimize the reconstruction loss. Studies have shown that AutoEncoders can effectively separate input data based on inherent patterns present in them[19, 40, 41].

To that end, we collected the embeddings of the training sequences from each respective PLM at different stages of their training. We trained an AutoEncoder using these final embeddings (Figure S2). We then projected the embeddings from different epochs, as shown in Figures 2b-d. Initially, the embeddings obtained for the different datasets were not mutually separated in the latent space (Figure 2b). This indicates that before training, the embeddings, regardless of the PLM used, exhibited similar patterns. We believe the embeddings at this stage are purely random, and thus appear alike to the AutoEncoder.

As training progressed and the models were trained, the embeddings began to reflect the differences in the molecular grammar of the datasets. This is evident as the embeddings of the three different datasets start to separate in the latent space (Figure 2c). Finally, upon further training, it is observed that each set of embeddings is distinctly separated in the latent space (Figure 2d). Therefore, we conclude that through training, the embeddings learned by different PLMs incorporate the molecular grammar of the sequences used for training.

### D. Statistical Interpretation of Protein Molecular Grammar

While we observe that the PLMs successfully capture the molecular grammar of the trained sequences, it remains in an abstract and unintelligible form to us. To gain insight into the probabilistic interpretation of the rules governing amino acid sequences, we construct the relative occurrence probabilities of a given amino acid next to another (either same or different) amino acid in a sequence (refer to Figures 3, and *SI Methods*).

**FIG. 3.**
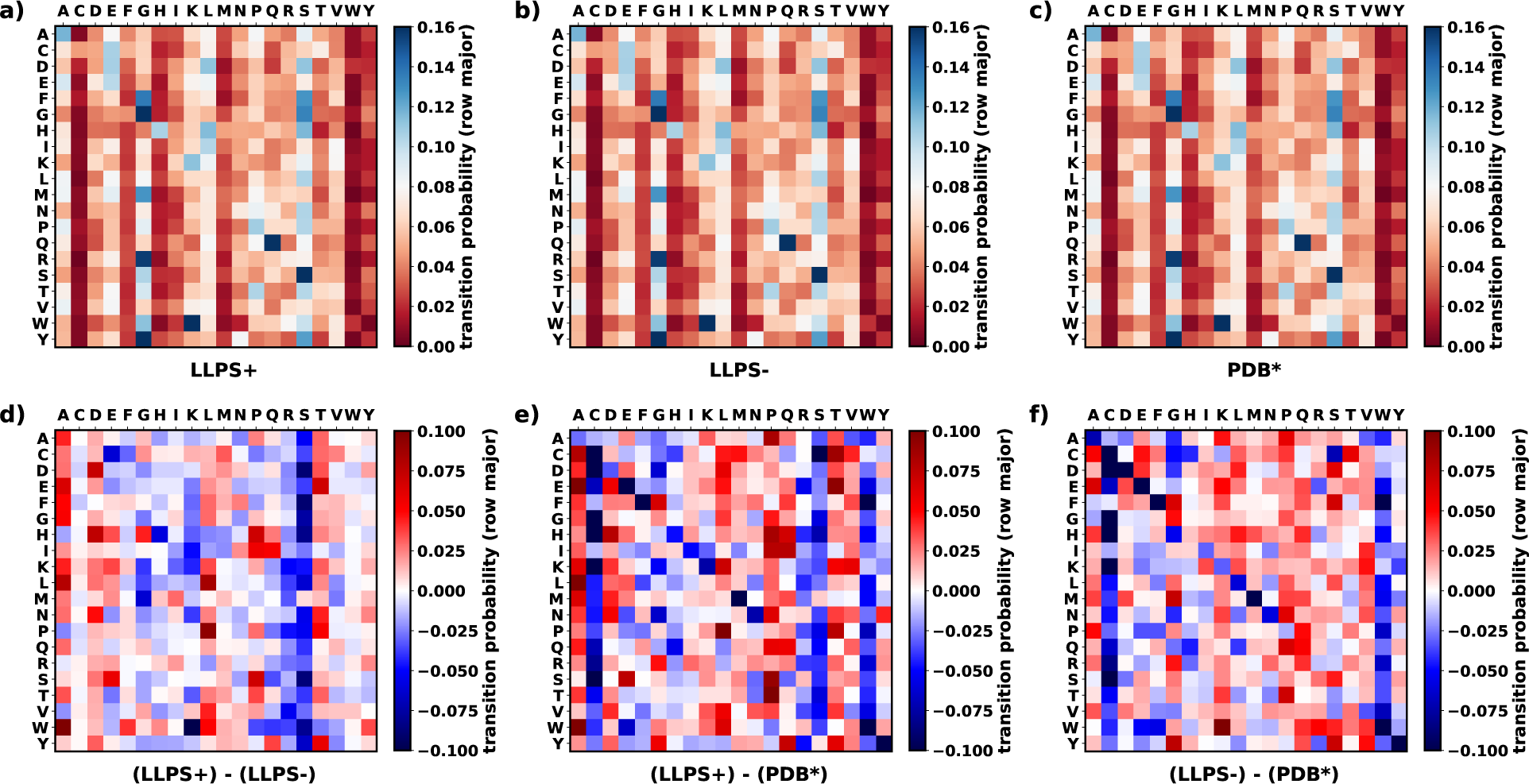
a-c) Heat map of relative occurrence probability among amino acids for lag=1 for the sequences generated from the GPT model of: **a)** LLPS+. **b)** LLPS-. **c)** PDB*. **d-e)** Difference of relative occurrence probability maps.

In Figures 3a-c it is evident that certain occurrence probabilities, specifically S-S, G-G, R-G, Y-H, W-K, and Q-Q, consistently exhibit high values regardless of the model as can be seen from the blue colour of those matrix elements. This means that in most of the sequences analyzed, one would find a serine followed by another serine, a glutamine by another glutamine and so on. These findings imply a universal trend across all proteins for these residue pairs, indicating a generic amino acid grammar. However, our primary interest lies in examining the distinctions in the relative occurrence probabilities of amino acids within the sequences generated by the three distinct GPT models.

Figure 3d illustrates the distinctions in transition probabilities between sequences generated by LLPS+ and LLPS-GPT models. An examination of the matrix reveals that a majority of the elements in the difference matrix between LLPS+ and LLPS-for datasets approach zero, indicating a high degree of similarity in their molecular grammar. Conversely, in Figures 3e, and 3f, there is a presence of dark red or dark blue colored elements, signaling that sequences generated by the PDB* GPT model exhibit a distinct molecular grammar from those produced by LLPS+ or LLPS-models. Notably, between LLPS+ and PDB*, we find that cysteine, serine, glycine and tryptophan have higher probabilities of appearing after any given amino acid in the LLPS+ sequences than the sequences belonging to PDB*. Apart from serine, the other three amino acids have higher probabilities of appearing in the LLPS-sequences as compared to PDB*. Therefore a major difference between LLPS+ and LLPS-sequences is the occurrence of serine after an amino acid in a sequence.

However, across all the difference plots (Figures 3d-f), the maximum disparity in transition probabilities remains low (around *∼*0.1). This suggests that the manifestation of Intrinsic Disordered Proteins (IDPs) does not arise from a drastic difference in their molecular grammar compared to that of folded proteins. Instead, it is these subtle differences that facilitate their ability to transition among multiple conformations, some or all of which may be aggregation-prone.

### E. Characteristics of the Predicted Protein Sequences

In Figure 4a, the distribution of sequence lengths for all generated sequences is depicted, with distinct colors corresponding to the model of origin. It is evident that there is no inherent bias discernible in the lengths of amino acid sequences generated across different models.

**FIG. 4.**
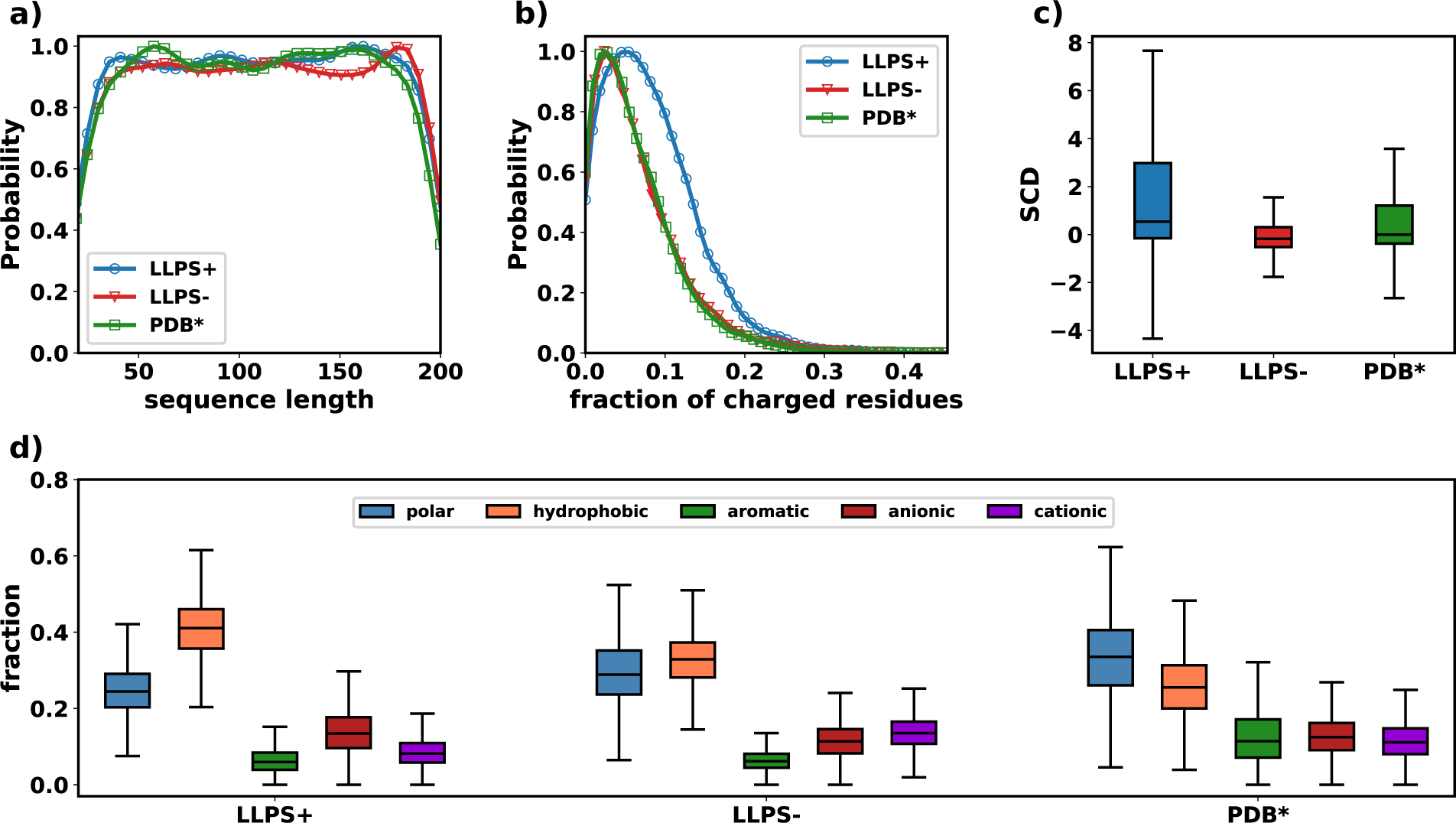
**a)** Distribution of sequence lengths for all three generated sequence sets. **b)** Distribution of the mean charge of a sequence for all three generated sequence sets. **c)** Comparison of sequence charge decoration (SCD) among the sequences generated from the different GPTs. **d)** Comparison of the probability of occurrence of amino acids in a sequence among the three generated sequence sets.

Subsequently, we scrutinized the fraction of charged residues (FCR) present in the sequences generated from each model (Figure 4b). It reveals a higher FCR for LLPS+ than LLPS- and PDB*. Additionally, they highlight the similarity in distributions of net and average charges between LLPS+ and PDB*. Notably, sequences generated via LLPS+ GPT exhibit the presence of more charged residues in a given sequence than the sequences generated from LLPS- or PDB*. This hints at the presence of extensive charge patterning in sequences from LLPS+ GPT.

We next calculated the sequence charge decoration (SCD) for all the sequences which are plotted in Figure 4c as per Eq-1. SCD determines the charge distribution within a given sequence, with higher SCD values potentially indicating a greater propensity for LLPS. The SCD of sequences predicted by the LLPS+ GPT model is higher compared to those generated by the LLPS-GPT and PDB* GPT models. Additionally, the SCD values from the sequences obtained from LLPS+ GPT model exhibit a broader distribution than those from the LLPS-GPT or PDB* GPT models. This suggests that the sequences predicted using the LLPS+ GPT model have a higher likelihood of forming LLPS, thus reinforcing confidence in the performance of our trained PLMs.

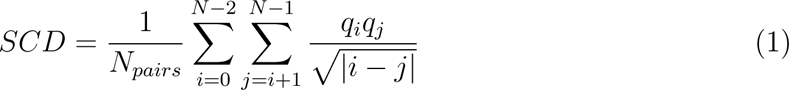

We next categorized the amino acid sequences into five types: polar, aromatic, hydrophobic, anionic and cationic. We then calculated the fraction of each type of residue per sequence for the generated sequences. We then plot their distributions via boxplots in Figure 4d. From Figure 4d, we observe that sequences generated from LLPS+ GPT have a higher fraction of polar residues (blue) than the rest. Aromatic residues (orange) seems to be low in all sequences in the dataset, which is a feature which the ML models learnt from the original dataset[27]. Recent investigations have also observed that replacing aromatic residues in a LLPS-prone sequence by non-atomatic residues does not preclude its phase separating ability[33]. We next observe that LLPS+ GPT sequences have a low fraction of hydrophobic residues while PDB* GPT sequences have the highest fraction of hydrophobic residues. The cationic and anionic residues seem to be present to the same extent in all sequences with a slight bias in the LLPS+ GPT sequences.

Overall, Figure 4 shows that sequences generated from LLPS+ GPT have higher amount of charged residues than the others, with a higher extent of charge patterning and a lower amount of hydrophobic residues. These strongly suggest that the sequences generated from LLPS+ GPT can potentially undergo phase-separation. However, simulations and experiments are required to confirm whether these sequences actually have phase separating potential or not.

### F. Experimental and Computational validation of LLPS formation potential in peptides from LLPS+ GPT

To validate whether the sequences predicted using LLPS+ GPT actually have a tendency to undergo LLPS, we performed computational and experimental validations. However, since the synthesis of full-length proteins is very complicated and tedious, we limited ourselves to short peptides of sequence lengths within 20 to 25 amino acids, all of which have been obtained from LLPS+ GPT. We generated 200 new sequences, with lengths between 20 and 25, using LLPS+ GPT. For each sequence the mean Low Complexity Region (LCR) probability was then calculated, which is the mean of the individual LCR forming probabilities of each amino acid in a single sequence. The mean pLCR is a probability that a given sequence has any conserved secondary structure or not. A higher value indicates that the sequence is mostly disordered and vice-versa. The LCR forming probabilities (pLCR) were predicted using the PrDOS webserver[42]. We selected 5 sequences from the 200 predicted sequences based on LCR probability. A summary of the selected sequences along with other sequence-based metrics have been provided in Table I.

**TABLE I.**
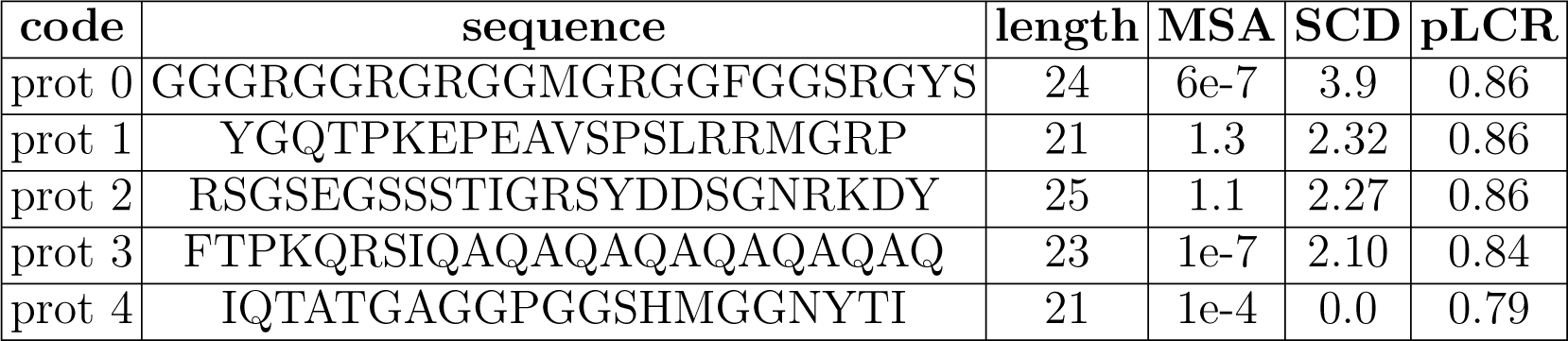
Summary of the peptides selected for experimental validation.

More importantly except for the prot0, all the other peptides show very low Multiple Sequence Alignment (MSA) when compared against the SwissProt database. This shows that except for prot0, all the other peptides are currently unknown. prot0 has the sequence **GGGRGGRGRGGMGRGGFGGSRGYS** which contains RGG repeats. RGG repeats are known to form homolytic LLPS[43]. Thus we use this as our positive control and perform validations initially on prot0.

#### 1. Validation of LLPS formation by the positive control (prot0)

We first tested the LLPS forming propensity using affordable computational methods. We conducted phase co-existence Molecular dynamics simulations (popularly known as ‘slab simulations’) at various temperature using a coarse-grained field namely CALVADOS2[44] that has been recently developed for IDPs (see *Methods* for simulation protocols). Using the mole fraction of the proteins present in the dense and the dilute phases, we plot the mol fractions vs the relative temperature of the simulations in Figure 5a to obtain a phase diagram.

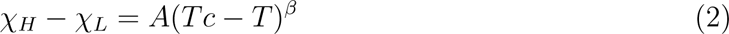

**FIG. 5.**
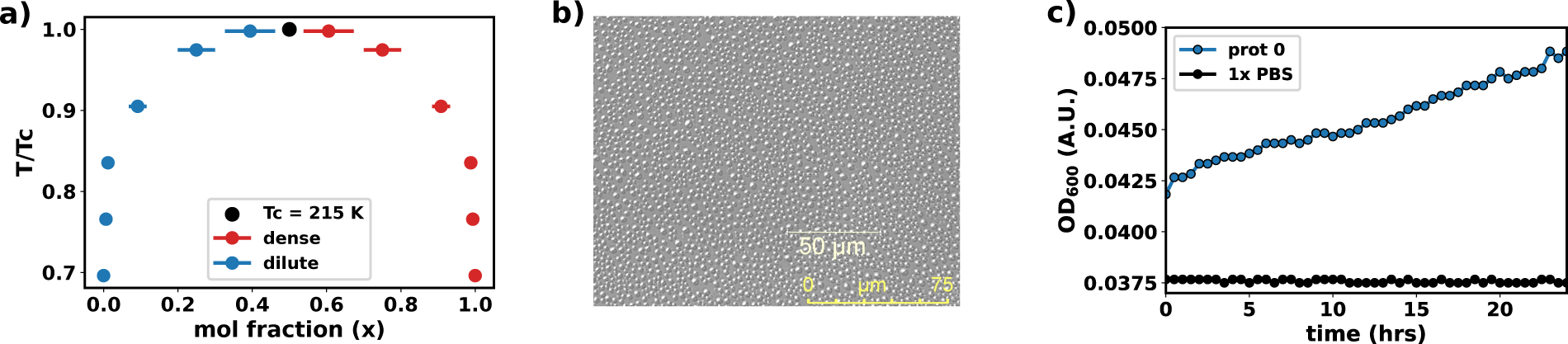
**a)** relative temperature vs. mole fraction phase-diagram for the predicted sequence. **b)** DIC microscopy image after incubation of 1mM of prot 0 in 1x PBS at 25 °C. **c)** Kinetics of droplet formation for prot0.

A fit of these points with Eq-2, we obtain *A*= 0.28 and *T_c_*= 215 K where *T_c_*is the upper critical solution temperature (UCST) and *A* is a parameter unique to the amino acid sequence. A value of *T_c_* = 215 K conveys that the generated sequence is predicted to undergo LLPS.

For a wetlab validation of simulation prediction, We then obtained the peptide in pure form (87%) from GenScript, New Jersey, USA (Figure S3a). A CD spectrum (Figure S3b) reveals that the peptide is completely disordered. A solution of 1 mM of prot 0 was prepared in 1x PBS (134 mM of Nacl at 7.4 pH) at 25°C and observed via a 20X air objective DIC microscopy (details in *SI Methods*). Figure 5b shows the occurrence of phase-separated droplets under the microscope. The kinetics of droplet formation (Figure 5c) shows a overall monotonic increase in droplets over a period of 24 hrs, explored via monitoring the absorbance at 600 nm. These conclusively prove the formation of LLPS by prot 0 under physiological conditions. Although the critical concentration of proteins used in training LLPS+ GPT are less than 100 *µ*M, we should take into account that prot 0 is a very short peptide while the proteins present LLPS+ dataset[27], which we used to train LLPS+ GPT, has an average length of 403*±*211 amino acids with a minimum length of 69 amino acids. Moreover, recent literature also shows that milimolar concentrations are generally used to investigate the LLPS by peptides[45–47]. These justify the usage of a higher concentration of the peptide during experiments. However, we have not investigated the critical concentration of prot 0 which can be lower than 1 mM.

#### 2. Formation of LLPS by the other selected peptides

Encouraged by the conclusive proof that prot 0 undergoes LLPS under ambient conditions, we obtained prot 1 and prot 2 in pure form (87%) from GenScript, New Jersey, USA (Figure S4-S5). However we could not obtain prot 3 with more than 70% purity (Figure S6). prot 4 was also obtained in the pure form (87%) from Galore TX, Bangalore, India (Figure S7). The CD spectra of prot1-4 reveal that all the peptides are completely disordered (Figure S8).

In 1x PBS at 25 °C and a concentration of 1 mM, pure Prot 1 and Prot 2, as well as semi-pure Prot 3, exhibit LLPS (Figure 6a-c, respectively). The kinetics of their droplet formation, shown in Figures 6d-e, further support their LLPS formation. The kinetics reveal that the LLPS of Prot 1 continues to grow over time. In contrast, the kinetics for Prot 2 show a quick saturation of OD_600_ at values lower than the OD_600_ values of prot 0, 1, or 3. This suggests a lower propensity for LLPS in Prot 2 compared to the others. However, Prot 4 does not show LLPS under these conditions. This indicates that its LLPS propensity, if present at all, is much lower than that of Prot 1-3.

**FIG. 6.**
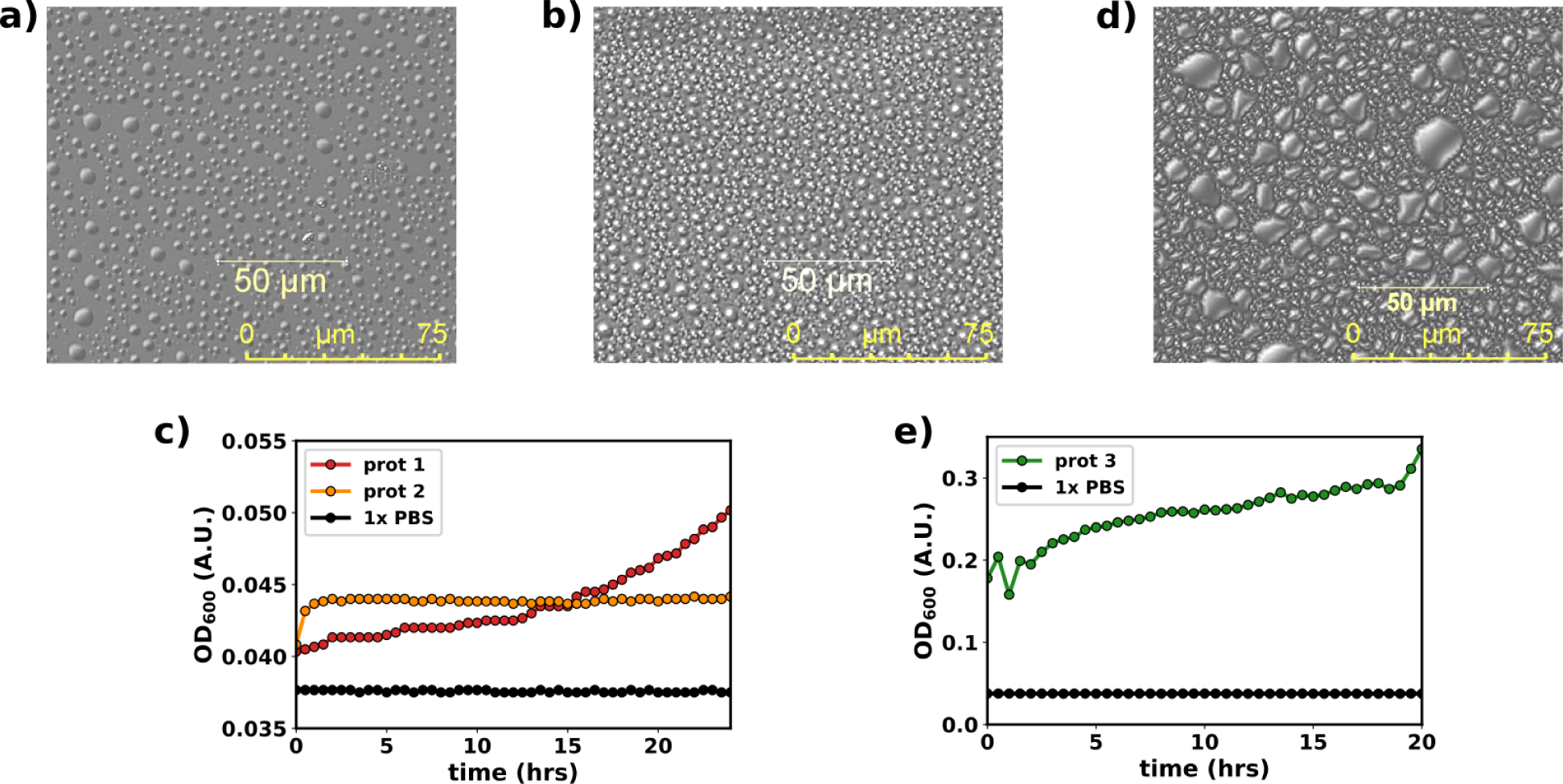
DIC microscopy image for the following peptides within 5 minutes of incubation in 1x PBS at 25 °C at 1 mM concentration: **a)** prot 1 (YGQTPKEPEAVSPSLRRMGRP). **b)** prot 2 (RSGSEGSSSTIGRSYDDSGNRKDY). **c)** prot 3 (FTPKQRSIQAQAQAQAQAQAQAQ). **d)** Kinetics for prot 1 and prot 2. **e)** Kinetics for prot 3.

Therefore we conclude that short peptides predicted using LLPS+ GPT have a high chance to be able to undergo LLPS in ambient conditions. Hence, we expect that full length proteins obtained from LLPS+ GPT will be highly prone to LLPS in ambient conditions. However conclusive proof will require further simulations and experiments which will be very tedious and should warrant a completely separate study altogether.

As a complimentary proof of concept on the successful generation of new sequences which will mimic the properties of the sequences used during the training of the PLM, we predicted a 67 residue protein using PDB* GPT.

**1 - HNCILYTEGCRWGLIPNMNLNDVRDWWRYWWP - 43**

**44 - MGPLLLHEWTSLKPPNWPLPEPESYTLLFHLEWYL - 67**

We obtained its possible structure using AlphaFold 2.0[48]. We observed that AlphaFold 2.0 predicts that the sequence in most likely to have a well folded structure (Figure S9a). Using this conformation as the starting point, we performed a 1 *µ*s molecular dynamics simulation (see *Methods* for details). Subsequent analysis of root mean squared deviation of the protein from the proposed AlphaFold 2.0 structure shows that the protein conformation does not change (Figure S9b) proving the fact that the generated sequence is that of a folded protein. This renders more credibility to the PLMs we have trained.

### G. Developing a Classifier to Differentiate LLPS-Prone and Non-LLPS Proteins

As discussed earlier, the number of known sequences that can undergo LLPS is extremely low in number. Thus, training a classifier would have not been possible due to either a very small dataset size or due to the large imbalance in the dataset with a dominant majority of it being composed of non-LLPS forming protein sequences. However, we have already generated synthetic sequences that can undergo LLPS as discussed in the previous sections. In an attempt to create a balanced dataset for training a classifier we use the 12000 sequences generated from LLPS+ and LLPS-GPTs as sequences that potentially have LLPS forming properties. We then extracted out the first 12000 protein sequences which do not contain any missing residues from the BMRB database (shared via Supporing file S4 in csv format). Combining these with the already present LLPS+, LLPS- and PDB* datasets[27], we obtained a total of about 24500 protein sequences that we used to train a classifier. We labelled sequences from the LLPS+ or LLPS-GPTs/dataset as 1, representing their potential to form LLPS. The sequences from the BMRB database or PDB* dataset were given a label of 0, i.e. that they cannot form LLPS.

However, the sequences are textual data and cannot directly be used in training. Therefore we used the recently reported ProstT5[37], an encoder-based PLM to obtain embeddings for the sequences. We used prostT5 to obtain vectorized representations of each amino acid for all the protein sequences followed by mean pooling along the amino acid sequence dimension to obtain a single vector representation of a fixed length (1024) for each sequence. We next prepared the training and testing datasets from the labelled, vectorized representations of the amino acid sequences as described in *Methods*. These were then used to train an e**x**treme **g**radient **b**oosted (XGB) classifier form the XGBoost python library[49] in a supervised manner (see *Methods* for details on hyper parameter tuning and model training).

We shuffled and split the 24500 sequences into two parts: 80% of it was used for training the classifier while the rest 20% was reserved for testing the classifier after post-training. After successful training, we made predictions on the sequences we had reserved for testing. The area under receiving operator characteristics (AUROC) curve (Figure 7a) shows a accuracy of 96% on the test dataset. The corresponding confusion matrix (Figure 7b) also shows that the classifier is also very accurate with a F1 score of 95%.

**FIG. 7.**
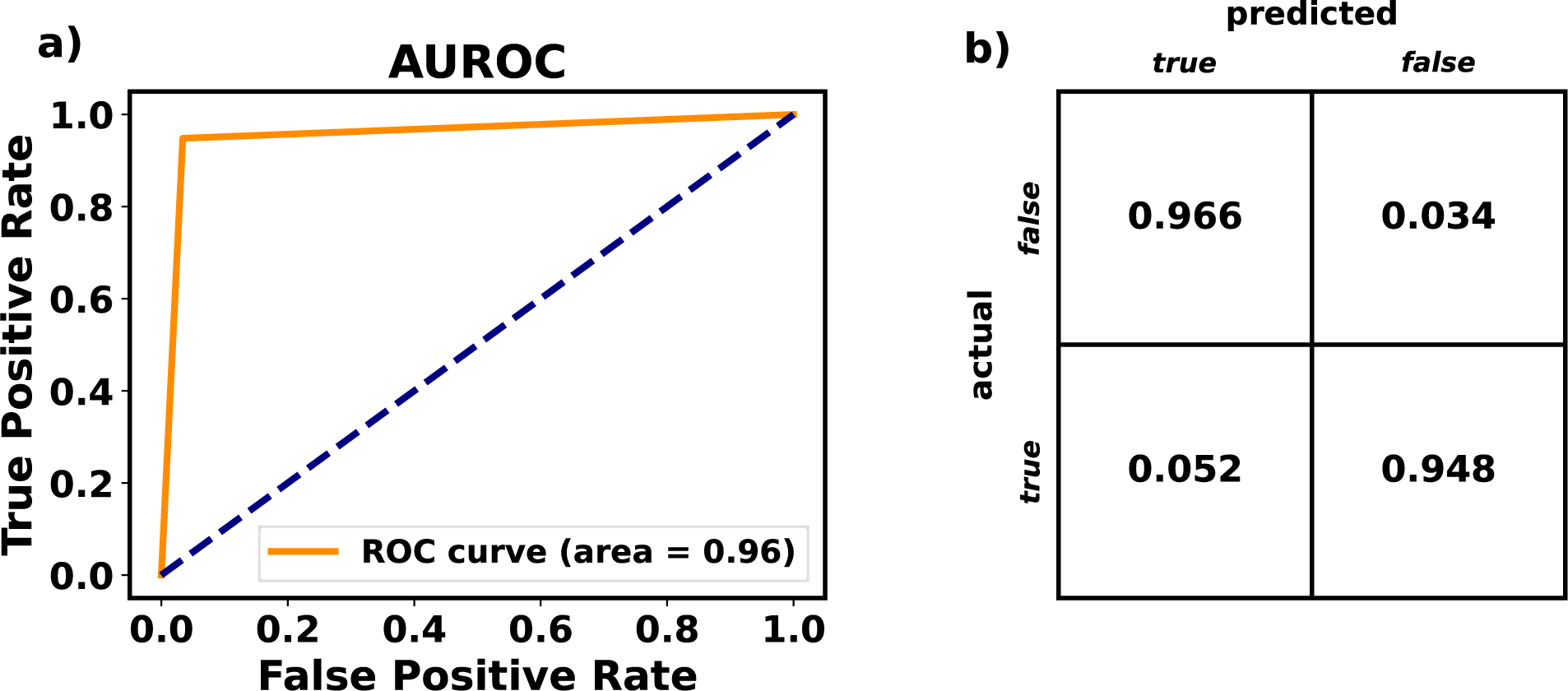
**a)** Area under receiving operator characteristics (AUROC) curve for the trained classifier. **b)** Confusion matrix for the classifier.

Therefore, with the help of the PLMs we trained, we were able to expand the dataset of LLPS forming proteins. Using these were then trained an XGB classifier which is very accurate in predicting whether a given sequence can undergo LLPS or not. To use the classifier developed, we have prepared a Google Colab notebook which can be accessed from https://bit.ly/4djlpgW.

## III. DISCUSSION AND CONCLUDING REMARK

In this investigation, we have developed and utilized transformer-based Protein Language Models (PLMs) to decode the molecular grammar inherent in proteins prone to liquid-liquid phase separation (LLPS). By training distinct PLM models on datasets representing varying LLPS propensities (LLPS+, LLPS-, and PDB*), we were able to generate three independent datasets each containing 6000 amino acid sequences. Our comprehensive investigation provided valuable insights into the probabilistic grammar that governs these sequences, shedding light on the subtle differences that facilitate phase separation in intrinsically disordered proteins (IDPs).

The generated sequences from LLPS+GPT and PDB*GPT demonstrated minimal re-semblance to known proteins in the SwissProt database, indicating the models’ capability to explore novel regions of the protein sequence space. The latent space analysis of sequence embeddings revealed distinct patterns, suggesting that our models effectively captured the unique characteristics of each dataset.

Statistical analysis of amino acid occurrence probabilities highlighted both universal trends and specific differences in the molecular grammar of IDPs and folded proteins. Experimental validation through computational and wet-lab methods confirmed the LLPS potential of small peptides derived from the LLPS+ GPT model. All atom MD simulation of a protein predicted by PDB* GPT showed that the protein has a rigid folded conformation. These support the reliability of our PLMs in predicting phase-separating or non-phase separating sequences.

The large embedding inherent in each of the PLMs encrypt the essence of “molecular grammar” that distinguishes the LLPS-prone GPT from that of LLPS-resistant GPT. It is the PLMs that have grasped the underlying rules. To gain insight into the “context” among amino acid pairs within a sequence, we extract attention maps for two representative sequences. Since the models have been trained to understand the molecular grammar inherent in an amino acid sequence and can predict the consecutive amino acids, given a sequence, the attention score among residues of a sequence would highlight the relationship among the residues that overall results in the properties of the sequence. Here the properties of interest are the secondary structure of a sequence and its ability to undergo homolytic LLPS. While grasping the connection of the propensity to undergo LLPS with attention scores is very difficult, the connection between attention scores and the secondary structure is simpler. High values of long-range attention scores (here a deeper blue hue) among residues hint at long range interactions among the residues leading to a well-defined secondary structure.

Figure 8a illustrates the attention scores derived from the LLPS+ GPT model for the initial 16 amino acids in FUS LCD (MASNDYTQQATQSYGA). Notably, besides the first row or column of the attention maps, instances of long-range contextual correlations are infrequent. Furthermore, there is no discernible pattern present in the sparse long-range correlations among amino acid pairs. In contrast, the attention scores obtained from PDB* GPT for the first 16 amino acids of Hen Egg White Lysozyme or HEWL (PDB ID: 1AKI) (KVFGRCELAAAMKRHG) (Figure 8b) immediately reveal significant contextual relationships among the initial amino acids. Additionally, higher contextual associations among amino acids farther from the sequence’s outset are evident compared to Figure 8a. We think that this is because HEWL possesses a highly rigid folded conformation, necessitating nearby amino acids to interact in a precise manner to uphold the three-dimensional fold. Conversely, for FUS LCD, the absence of a specific residue-residue “contexts” contributes to its identity as an Intrinsically Disordered Protein (IDP), as it does not have a rigid fold. In summary, our observations indicate that IDPs lack long-range contextual information among amino acids, while folded proteins likely exhibit more long-range context among amino acid pairs, essential for maintaining their conformation and function.

**FIG. 8.**
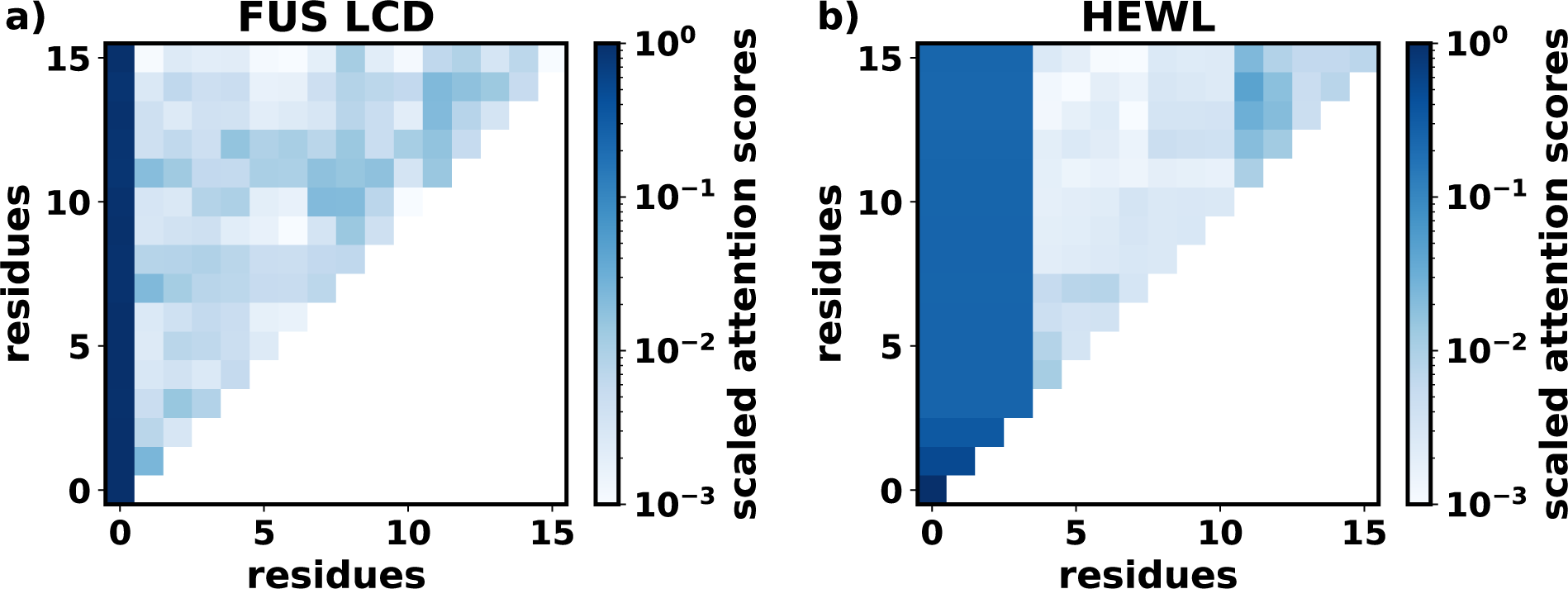
**a)** Attention map for the first 16 residues of FUS-LCD. **b)** Attention map for the first 32 residues of HEWL (PDB ID:1AKI).

In the last part of the investigation, we report the development of a robust classifier using sequences generated by our PLMs, achieving high accuracy in distinguishing LLPS-prone proteins from non-LLPS proteins. Our approach acknowledges that while LLPS-forming propensity is embedded in a protein’s primary structure, current metrics and models are too simplistic to capture the extremely complex grammar present in such sequences. From our observations in this study and previous research [18, 33, 47], it appears that none of the current mainstream metrics[44, 50] are particularly sensitive. This hypothesis is further supported by recent studies [33, 47]. In light of this, machine-learnt classifier such as the one developed here, can do a better job than various metrics (many of which has been investigated in the present investigation) for characterizing and distinguish LLPS-prone sequences from non-LLPS sequences. By utilizing transformer-based models, we aim to better understand and predict the molecular grammar governing LLPS, providing a more robust and sensitive method for identifying LLPS-prone sequences.

## IV. METHODS

### A. Training datasets for PLMs

In this investigation, we utilized the already reported LLPS+, LLPS-, and PDB* datasets[27].

The LLPS+ dataset encompasses 137 amino acid sequences of intrinsically disordered proteins (IDPs) with critical/saturation concentrations either identical or lower than 100 µM. In contrast, the LLPS-dataset comprises 84 amino acid sequences of IDPs with critical/saturation concentrations exceeding 100 µM.

The PDB* dataset consists of 1562 folded proteins that do not spontaneously undergo liquid-liquid phase separation (LLPS) unless under extremely drastic conditions.

Protein sequences were denoted using the single letter codes for amino acids. Notably, all datasets include proteins with seleno-cysteine, denoted by the single-letter code U, and proteins with missing residues, represented by the letter X. To prepare the protein sequences of each dataset for training, a single text file was generated for each dataset. Within these files, protein sequences were separated by a “blank space” to indicate the end of a sequence.

We used a character level tokenization for training resulting in a vocabulary of the following characters: **ACDEFGHIKLMNPQRSTUVWXY***<***blank space***>*.

### B. Tokenization and positional embeddding

For any PLM to work, we need to convert the characters to numbers that the machine can understand. Thus we use character-level tokenization (*t*) of the amino acid sequences to convert a protein’s primary sequence to an array of numbers, more specifically integers. The character level tokenization has been summarized in Table-II

**TABLE II.** The numeric representation of the vocabulary.

| character | A | C | D | E | F | G | H | I | K | L | M | N | P | Q | R | S | T | U | V | W | X | Y | <blank space> |
| --- | --- | --- | --- | --- | --- | --- | --- | --- | --- | --- | --- | --- | --- | --- | --- | --- | --- | --- | --- | --- | --- | --- | --- |
| token | 1 | 2 | 3 | 4 | 5 | 6 | 7 | 8 | 9 | 10 | 11 | 12 | 13 | 14 | 15 | 16 | 17 | 18 | 19 | 20 | 21 | 22 | 23 |

To comprehend the context or significance of the sequence order, Protein Language Models (PLMs) require information about the position of each token within a given sequence. Achieving this involves the use of positional embedding. The positional embedding value for any character at position *p* in a sequence of length *l* is determined by Eq-3[34].

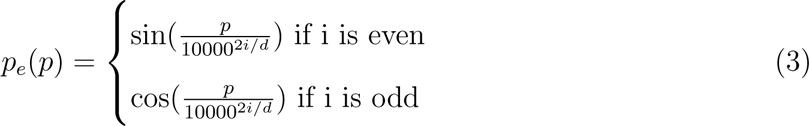

where *i* is the positional encoding along the *i*-th dimension for a *d* dimensional embedding of the tokens.

Therefore, the final numerical representation of a character in a given sequence is obtained by adding the token embedding and the positional embedding. Through this process, sequences of characters are transformed into machine-understandable numerical values, while retaining crucial information about the sequential order of the characters.

### C. GPT model details

We employ a decoder-only GPT model with an architecture resembling Figure-1a). Specifically designed for amino acid sequences, this model serves as an advanced tool for sequence generation, eliminating the need for an encoder component. Trained on a specific type of protein sequences, the model excels at predicting subsequent amino acids given an initial sequence or amino acid, ensuring that the generated sequence reflects the properties of the proteins used during training.

In essence, the model undergoes a comprehensive training phase to grasp the intricate language of amino acids, analogous to mastering the grammatical structure of a language. When presented with a partial protein sequence of length *l* as input, the model leverages its acquired knowledge to analyze the context of surrounding amino acids. This analysis is facilitated by multiple layers (*N*) of transformer neural networks, each utilizing a “multiheaded self-attention” mechanism to comprehend the contextual nuances.

Our current implementation of the decoder-only GPT model features 8 layers of multiheaded self-attention (*N* = 8), with each layer containing 8 heads (*N_h_* = 8). The chosen sequence length for training is *l* = 128. This meticulous configuration optimizes the model’s performance across diverse protein sequences.

The core process involves generating a probability distribution for potential amino acids at the next position in the sequence based on the learned context. The model then selects the most probable amino acid, iteratively extending the sequence. Noteworthy is the model’s adaptability, enabling it to provide diverse sequence predictions and offer multiple potential outcomes instead of adhering to a rigid pattern.

The term “pre-trained” emphasizes the model’s wealth of knowledge acquired during its initial training on a dataset as vast as possible. This foundational knowledge allows the model to discern general protein sequence patterns, facilitating the generation of meaningful and contextually relevant sequences even with partial input. The model converts partial input into an abstract vector representation with a length of *d*, subsequently transforming it into probabilities from which the next amino acid is predicted. For our study, we set *d* = 256.

### D. Training details for PLMs

The training process for each model in our study involved a substantial training duration of 10^6^ epochs, ensuring a thorough exploration of the model’s learning capabilities. We adopted a batch size of 64 to efficiently process and update the model parameters. The choice of the cross-entropy loss function, a standard measure for evaluating the dissimilarity between predicted and actual sequences, guided the training process. Additionally, we employed the AdamW optimizer with a specified learning rate of 0.0001. AdamW, renowned for its effective weight decay mechanism, contributed to the optimization of model parameters during training.

To assess the model’s performance and generalization, we meticulously monitored both training and validation losses. The evolving trends in these losses over the epochs provide crucial insights into the model’s learning dynamics and its ability to adapt to new data. Detailed visualizations of the training and validation losses are presented in Figure-S1, offering a comprehensive understanding of the model’s progression. The chosen metrics and optimizer configuration aim to strike a balance between model convergence and preventing overfitting, ensuring that the trained models generalize well to new, unseen data. Overall, this rigorous training regimen, coupled with systematic loss monitoring, establishes a robust foundation for the subsequent evaluation and analysis of the model’s predictive capabilities.

### E. Generation of sequences from the PLMs

We initiated the generation of 6000 sequences for each of the GPT models: LLPS+, LLPS-, and PDB* (and hence total 18000), as described in the *Results* section. These sequences, ranging in length from 20 to 200, were initialized with a random character from the vocabulary. It’s noteworthy that the vocabulary includes X, U, and *<*blank space*>* as characters. To ensure the correctness of the predicted sequences, we made slight adjustments:

1. Any occurrence of *<*blank space*>* was removed.
2. Subsequently, all X’s, if present, were removed.
3. Finally, any instance of U was replaced with C.

However, as discussed in the *Results* section, at most *l_max_*(= 128) amino acids can be passed into the model at a time. To generate sequences longer than *l_max_* + 1, one must use the last *l_max_* amino acids as input. The model’s output can be appended to the last *l_max_ −* 1 amino acids of the input sequence, thus maintaining the sequence length at *l_max_*, which can then be fed back into the model.

### F. All-atom molecular dynamics simulation details

GROMACS-2023.2 was employed for conducting all-atomistic molecular dynamics simulations[51, 52].The charmm36m forcefield[53] and the charmm modified TIP3P water model[54] was used to simulate folded structures. Long-range electrostatics were calculated using Particle Mesh Ewald (PME)[55] summation, while Lennard–Jones interactions were determined using the Verlet cutoff scheme[56], with 1.2 nm as the cutoff for both. The LINCS[57] algorithm was employed to constrain hydrogen bonds, and the SETTLE[58] algorithm was used to fix hydrogen bonds in water molecules. Initial steps involved solvating the simulation system, followed by achieving electro-neutrality by introducing Na^+^ and Cl*^−^* ions. Additional 100 mM NaCl ions were incorporated to maintain physiological concentrations of NaCl throughout the simulations.

Subsequently, the system underwent energy minimization, followed by equilibration in an NVT ensemble for 100 ps, employing a timestep of 2 fs. This was succeeded by equilibration in an NPT ensemble for 100 ps, utilizing a timestep of 2 fs. All simulations were conducted at 310.15 K, with a reference pressure of 1 bar, where applicable. To control temperature and pressures, the v-rescale thermostat[59] and the c-rescale barostat[60] were employed. The production runs were then executed using the respective equilibrated systems, with a timestep of 2 fs.

### G. Phase co-existence molecular dynamics simulation details

Phase coexistence molecular dynamics were performed using CALVADOS 2[44]. For the generation of the initial conformations, we used the software “packmol”[61] and packed 100 protein chains inside a cubic box with a side length of 30 nm. We then performed NVT simulations at a low temperature (120.15 K) for 1 *µ*s with a timestep of 10 fs using OpenMM[62]. This resulted in a single droplet being formed.

Starting with the final conformation obtained from the simulation at the low temperature (120.15 K), we performed simulations at different higher temperatures. The Langevin integrator with a friction coefficient of 1.0 ps*^−^*^1^ and a time step of 10 fs was used to perform NVT simulations. An ionic concentration equivalent to 150 mM of NaCl was maintained throughout the simulations. These simulation parameters were adapted from [44]. The simulation frames were saved at 100 ps intervals. The last 1 *µ*s of each trajectory was used for calculation of the phase diagram.

### H. Materials

The peptides prot 0, prot 1 and prot 2 were purchased from GenScript, New Jersey, USA with purity *≥* 87%, with N- and C-terminal acetylated and amidated, respectively. prot 3 was chemically synthesized via solid phase peptide sythesis (SPPS) by Anirban and Ushasi, again with N- and C-terminal acetylated and amidated, respectively. However, it was possible to obtain prot 3 at not more than 70% purity. prot 4 was purchased from Galore TX, Bengaluru, India. The mass of each peptide was confirmed by MALDI-TOF-MS peptide mass analysis (Shimadzu MALDI-8030; with 500 scans for each run having laser power as 20 mV), using *α*-cyno-4-hydroxycinnamic acid matrix (Bruker Daltonics, MA).

### I. Turbidity assay

The absorbance of 1 mM peptide solutions (1x PBS buffer, pH 7.4) was measured at *λ*=600 nm using BioTek Synergy H1 Microplate Reader, kinetically over a period of 24 h. The readings were taken at regular intervals of 30 min with 5 sec shaking (orbital) before each measurement. Each data represents biological replicates of two days with n=3.

### J. DIC microscopy

The phase separation behavior of 1 mM peptides in 1x PBS was studied visually using NCI Leica DMC4500 Microscope, in the reflection mode, with differential interference contrast (DIC) with 20x optical zoom lens. Images were taken with a DMC4500 camera under the control of LEICA LAS X software (Version 3.4.2).

### K. CD spectra measurement

The CD spectra of the peptide solutions were recorded on Jasco J-815 Circular Dichroism Spectrometer. The spectra (average of 3 scans for each sample) were collected within the wavelength range of 200 nm and 260 nm having a bandwidth of 1.00 nm, 0.5 nm step. Solvent (1x PBS) background was subtracted from each spectrum of 0.25 mM of each peptide spectra.

### L. Dataset preparation for the LLPS classifier

All the sequences (generated + real) were first pooled and subsequently labelled as LLPS forming (1) or not LLPS forming (0). This resulted in a total of 24500 sequences. The amino acid sequences were converted into vector representations by using prostT5[37], an encoder only PLM that has been pre-trained on 2.2 M protein sequences present in AlphaFold DB. Hence, we can assume that prostT5 has learnt the generic grammar of proteins can be safely used to obtain context-based embeddings for a given sequence of amino acids. Therefore, for a sequence of length *l*, the output representation was a tensor of the shape (*l,* 1024). A Mean Pooling was then performed along the sequence length dimension to obtain a single vector, 1024 long. Thus each sequence could be represented by a single vector of a fixed length. The labelled vectors were then generously shuffled and 20% of them were separated for testing. The rest 80% were used to train an e**x**treme **g**radient **b**oosted (XGB) classifier[49].

The primary task was to determine the optimum hyper parameters for the classifier. For accurate determination of the this, we used the GridSearchCV function from python’s scikit-learn[63, 64] library and obtained an optimum value of maximum tree depth = 4 and number of estimators = 1024 for the classifier given it is being trained on the dataset we designed.

### M. List of software

We have used only open-source software for this study. All simulations have been performed using GROMACS-2021[51, 52]. Snapshots were generated using PyMOL 2.5.4[65–67]. Analysis were performed using Python[68] and MDAnalysis[69, 70]. Figures were prepared using Matplotlib[71], Jupyter[72] and Inkscape[73]. All deep neural network models were built using PyTorch[74]

## V. DATA AND CODE AVAILABILITY

The scripts and models used to train GPT models and predict sequences have been uploaded to https://github.com/JMLab-tifrh/idpGPT. All relevant data have been uploaded to a Zenodo repository (https://zenodo.org/records/13293185). For easy usage of the PLMs to generate new sequences, we have created a Google Colab notebook which is available at https://bit.ly/4clObMP. Similarly, to classify sequences using the trained XGB classifier, another Google Colab notebook has been prepared which is available at https://bit.ly/4djlpgW.

## ACKNOWLEDGEMENT

We would like to thank Prof. Anand Srivastava and Mr. Santosh Prajapati for their insights in performing on phase-coexistence simulations and useful discussions. We thank Dr. Aneesh Tazhe Veetil for insights on the experiments. We also thank Prof. Kalyaneshwar Mandal and Mr. Anandhmuthiah V for useful discussions regarding the LLPS propensities of the generated sequences. We acknowledge support of the Department of Atomic Energy, Government of India, under Project Identification No. RTI 4007. JM acknowledges Core Research grants provided by the Department of Science and Technology (DST) of India (CRG/2023/001426).

