## Supplemental figures and tables for "Harnessing Transformers to Generate Protein Sequences Prone to Liquid Liquid Phase Separation"

#### for

### I. SUPPLEMENTARY METHODS

#### A. Calculation of occurrence probability matrices

Occurrence probabilities between amino acids within protein sequences were computed using a custom Python algorithm implemented in this study. For each protein sequence in the dataset, the algorithm iterates over the sequence, accumulating occurrence counts between amino acids with a lag of 1 in a occurrence count matrix $T$, that is it counts occurrences between adjacent amino acids.

Additionally, the number of occurrences from an amino acid counted in the count vector $C$. occurrence probabilities are then computed by normalizing the occurrence count matrix by the total count of each amino acid, ensuring that probabilities sum up to 1 along each row. This is represented by the Eq-1:

$$P(i,j) = \frac{T(i,j)}{C(i)} \tag{1}$$

To prevent division by zero errors, amino acids with zero occurrences are assigned a count of 1 before computing occurrence probabilities.

#### B. Details of the autoencoder architecture

We employed an autoencoder architecture to distill protein sequence embeddings into a succinct two-dimensional latent space, enabling effective dimensionality reduction and feature extraction. Our autoencoder comprises an encoder and decoder, each composed of multiple dense layers. The encoder initiates with an initial dense block and progresses through a sequence of dense layers, gradually reducing input dimensionality. Specifically, we adopted a layer configuration of [1024, 256, 64, 32, 8] for the encoder, progressively decreasing layer sizes. This approach facilitates hierarchical feature extraction from the input protein sequence embeddings. Subsequently, a latent layer compresses the representation into a two-dimensional space, capturing essential input characteristics.

Conversely, the decoder mirrors the encoder's architecture in reverse, endeavoring to reconstruct the original input from the compressed latent representation. The decoder also incorporates multiple dense layers, with dimensions symmetrically increasing towards the output layer. During training, we employed the Mean Squared Error (MSE) loss function and utilized the Adam optimizer with a learning rate of $1 \times 10^4$ to minimize reconstruction error and iteratively update model parameters.

We extracted embeddings for each amino acid from respective protein sequences using PLMs, resulting in vector representations of length 256 for each amino acid. Thus, for a sequence of length $l$, the corresponding representation obtained was $(l, 256)$. The sequence's vector representation was then obtained by Mean Pooling the vectors along the sequence length. Subsequently, we utilized these embeddings to train the autoencoder, projecting the embeddings onto the latent space. Ultimately, we obtained a two-dimensional latent space, as detailed in the main text, for further analysis and interpretation.

#### II. SUPPLEMENTARY FIGURES

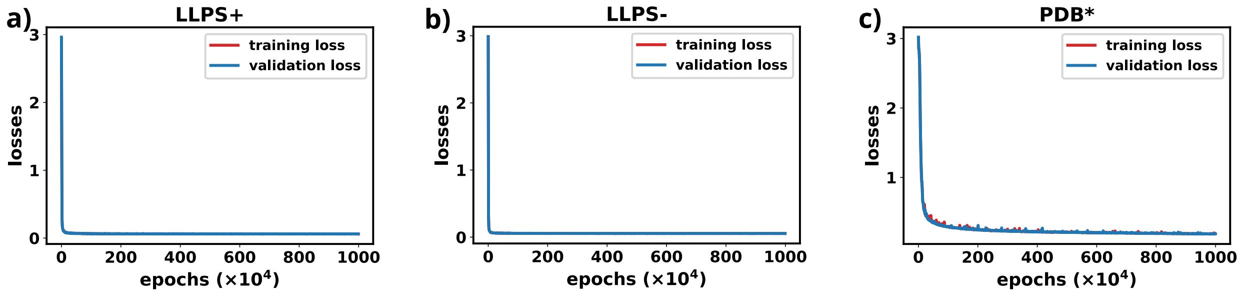


FIG. S1. Training and validation loss for **a)** LLPS+ GPT. **b)** LLPS- GPT .**c)** PDB* GPT

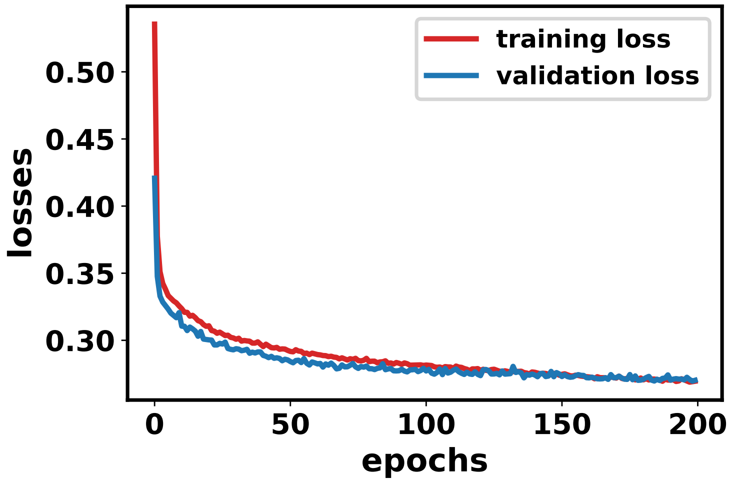


FIG. S2. Training and validation losses of the AutoEncoder.

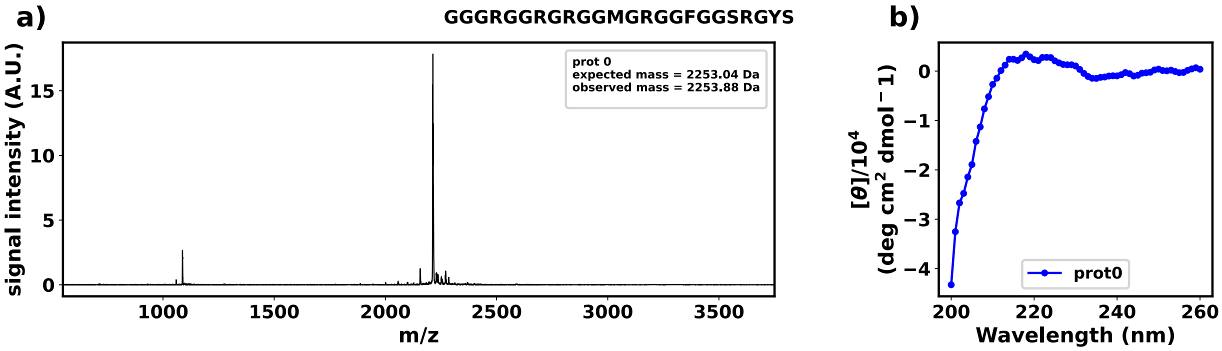


FIG. S3. **a)** Mass spectroscopy data for prot 0. **b)** Circular dichroism spectroscopy for prot 0.

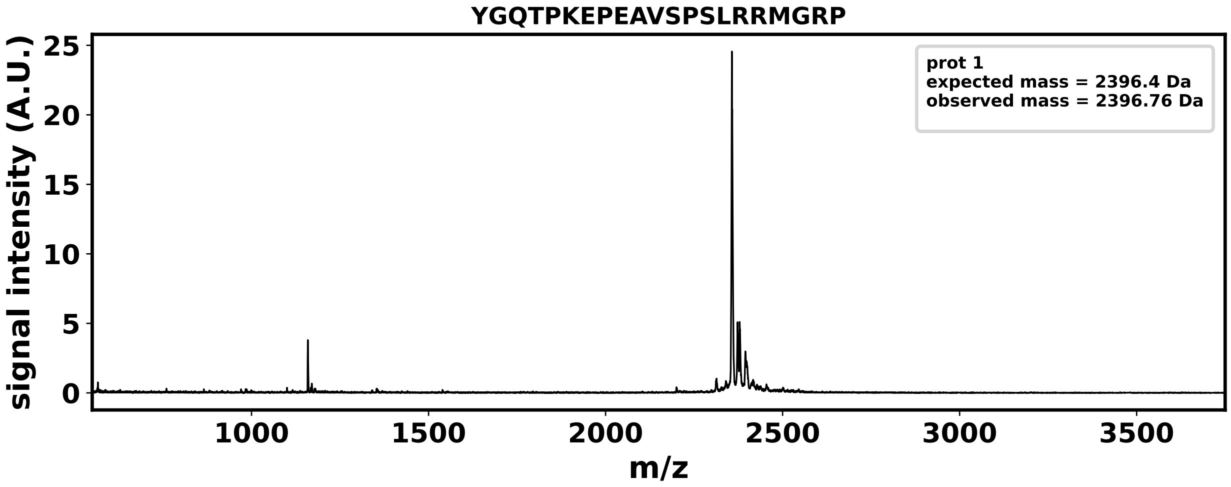


FIG. S4. Mass spectroscopy data for prot 1.

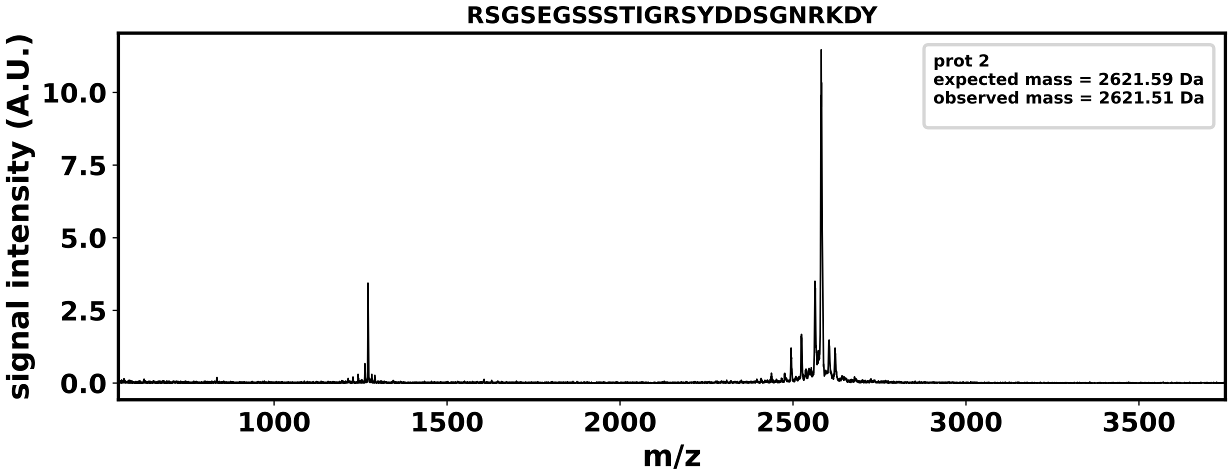


FIG. S5. Mass spectroscopy data for prot 2.

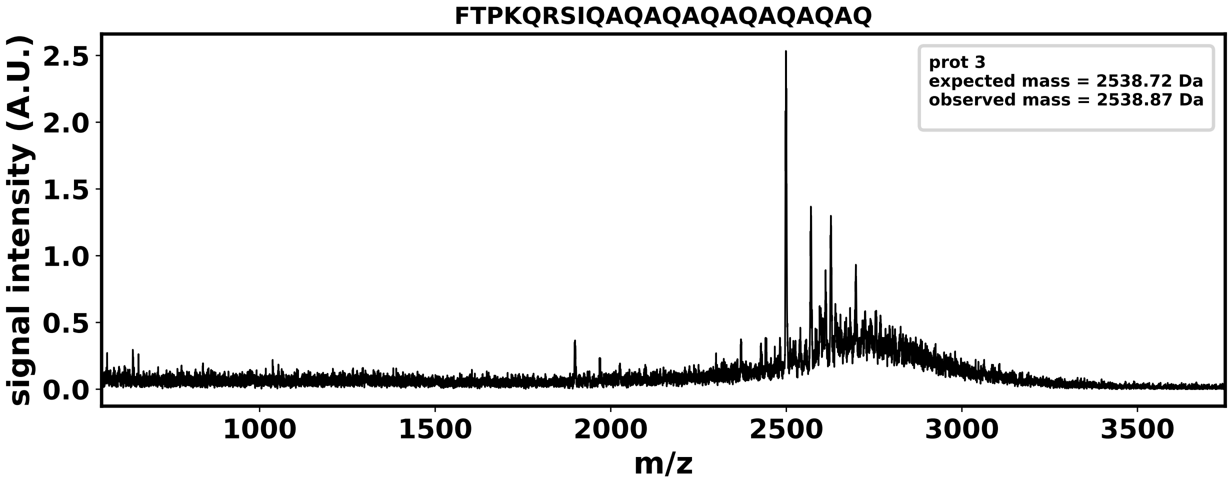


FIG. S6. Mass spectroscopy data for prot 3.

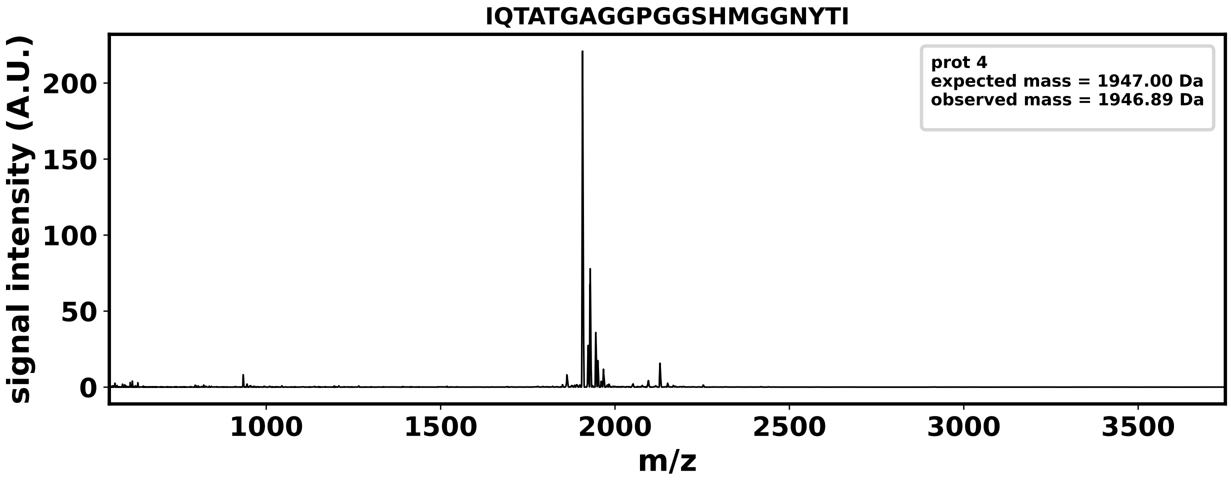


FIG. S7. Mass spectroscopy data for prot 4.

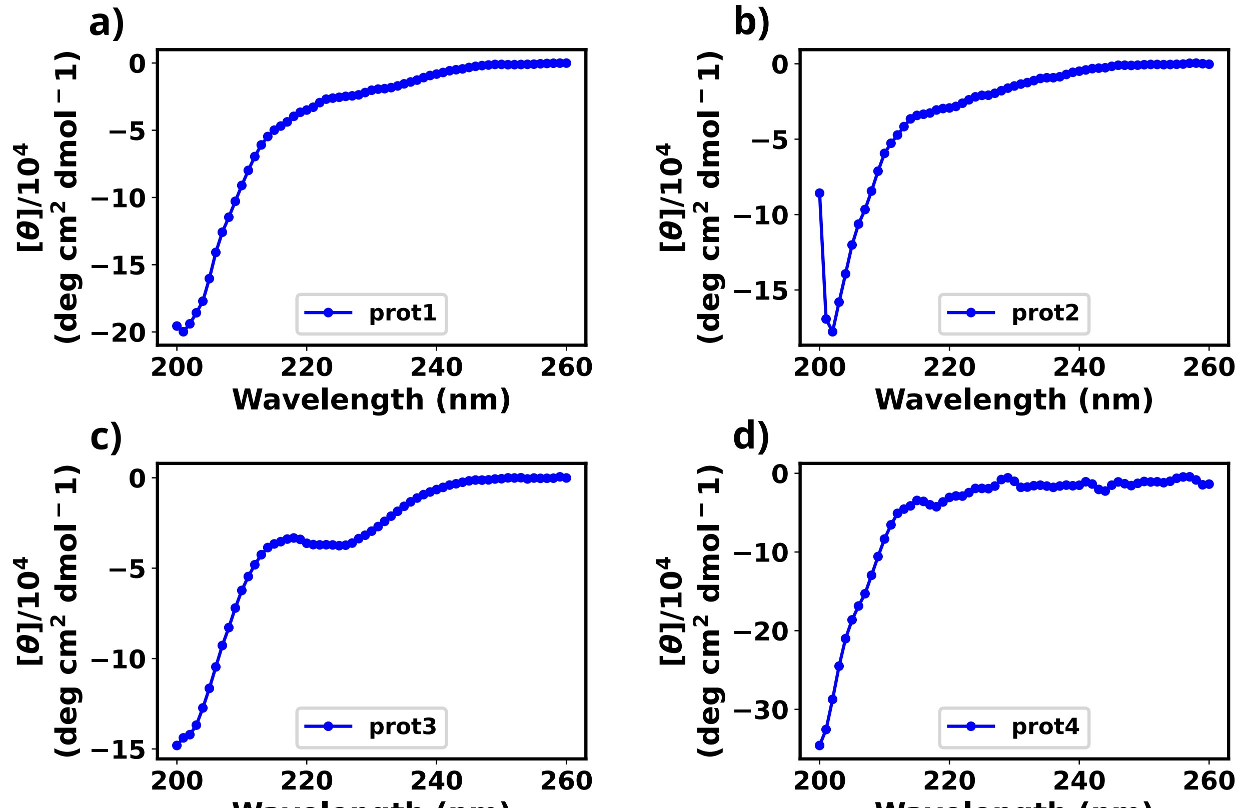


FIG. S8. Circular dichroism spectra for **a)** prot 1. **b)** prot 2. **c)** prot 3. **d)** prot 4

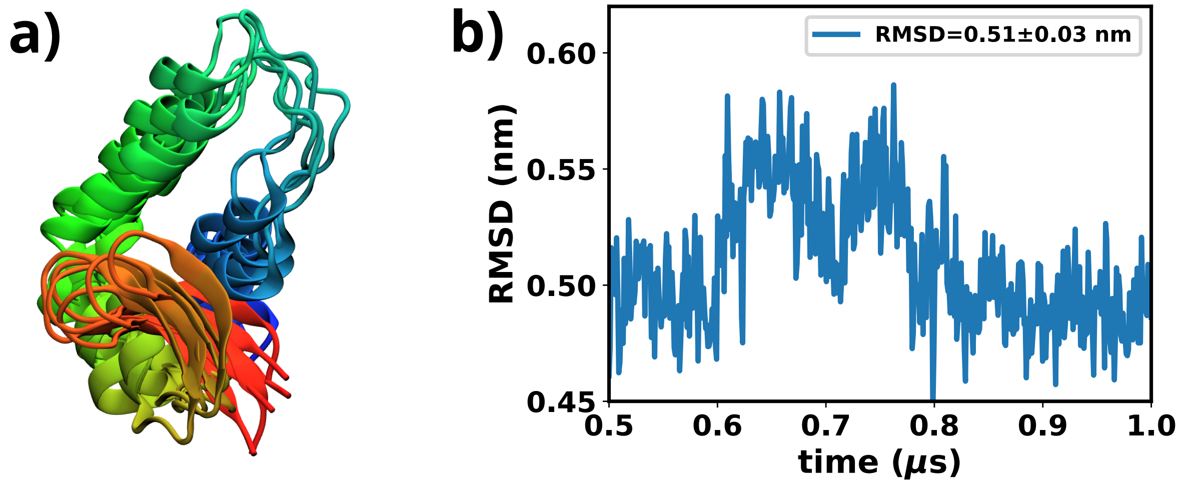


FIG. S9. **a)** Overlap of 5 conformations obtained from the last 500 ns of simulations. The conformations are 100 ns apart from each other. **b)** Time profile of RMSD for the last 500 ns for the folded parts of the protein.
